# SpyDisplay: A Versatile Phage Display Selection System using SpyTag/SpyCatcher Technology

**DOI:** 10.1101/2022.12.19.520987

**Authors:** Sarah-Jane Kellmann, Christian Hentrich, Mateusz Putyrski, Hanh Hanuschka, Manuel Cavada, Achim Knappik, Francisco Ylera

## Abstract

Phage display is an established method for the *in vitro* selection of recombinant antibodies and other proteins or peptides from gene libraries. Here we describe SpyDisplay, a phage display method in which the display is achieved via SpyTag/SpyCatcher protein ligation instead of genetically fusing the displayed protein to a phage coat protein. In our implementation, SpyTagged Fab antibody fragments are displayed via protein ligation on filamentous phages carrying SpyCatcher fused to the pIII coat protein. A library of genes encoding Fab antibodies was cloned in an expression vector containing f1 replication origin, and SpyCatcher-pIII was separately expressed from a genomic locus in engineered *E. coli*. We demonstrate the functional, covalent display of Fab on phage, and rapidly isolate specific high-affinity clones via phage panning, confirming the robustness of this selection system. SpyTagged Fabs – the direct outcome of the panning campaign - are compatible with modular antibody assembly using prefabricated SpyCatcher modules and can be directly tested in diverse assays. Furthermore, SpyDisplay streamlines additional applications that have traditionally been challenging for phage display: we show that it can be applied to N-terminal display of the protein of interest and it also enables display of cytoplasmically folding proteins exported to periplasm via the TAT pathway.

## Introduction

The *in vitro* selection of peptides and proteins with desired properties from large gene libraries (Jijakli et al., 2016) is a powerful approach that has been used extensively for the discovery of binding molecules (Jijakli et al., 2016), including antibodies (Winter et al., 1994). All *in vitro* selection technologies require the physical linkage of genotype and phenotype. Phage display is the oldest and most widely used method due to its robustness and favorable properties such as speed, simplicity, and accommodation of large libraries. To couple genotype to phenotype, the proteins to be selected are displayed on the surface of engineered filamentous M13 phages, which in addition contain the genetic information of the presented proteins. In conventional phage display, the displayed protein is genetically fused to a coat protein of the phage, in most cases to the minor coat protein pIII. Instead of a full phage genome, smaller plasmid derivatives called phagemids are commonly used in phage display, as they simplify cloning and allow the creation of larger libraries (Breitling et al., 1991; Qi et al., 2012). Such phagemids contain the genes for the displayed proteins fused to the gene encoding the phage coat protein used for display, antibiotic resistance genes, and genetic elements required for plasmid-like replication as well as for replication of ssDNA and its packaging in phage capsid. Phage display with phagemids necessitates the use of helper phage, which provides the remaining structural and regulatory phage proteins that are not present in the phagemid and thus allows phage assembly after superinfection of phagemid-containing bacteria. After several rounds of phage display, the selected genes are typically subcloned into a suitable expression plasmid (Dubel et al., 1993) for screening and further analysis.

The SpyTag/SpyCatcher protein ligation technology (Zakeri et al., 2012) is a versatile method to covalently link two proteins. The SpyTag, a short peptide of 13 amino acids, reacts spontaneously with SpyCatcher protein (12.3 kDa) to form an isopeptide bond between an aspartic acid residue in the tag and a lysine residue in the Catcher, crosslinking the two (Fig. 1A: SpyTag/SpyCatcher system). The reaction is fast, specific and has been further optimized in form of the SpyTag2/SpyCatcher2 (Keeble et al., 2017) and SpyTag3/SpyCatcher3 (Keeble et al., 2019) systems. This technology has been used in many different applications, for example in the production of vaccine nanoparticles or stabilized enzymes (Keeble and Howarth, 2020). Furthermore, it has recently been applied for modular antibody assembly and site-specific labeling of antibodies (Alam et al., 2017; Hentrich et al., 2021).

**Figure 1:**
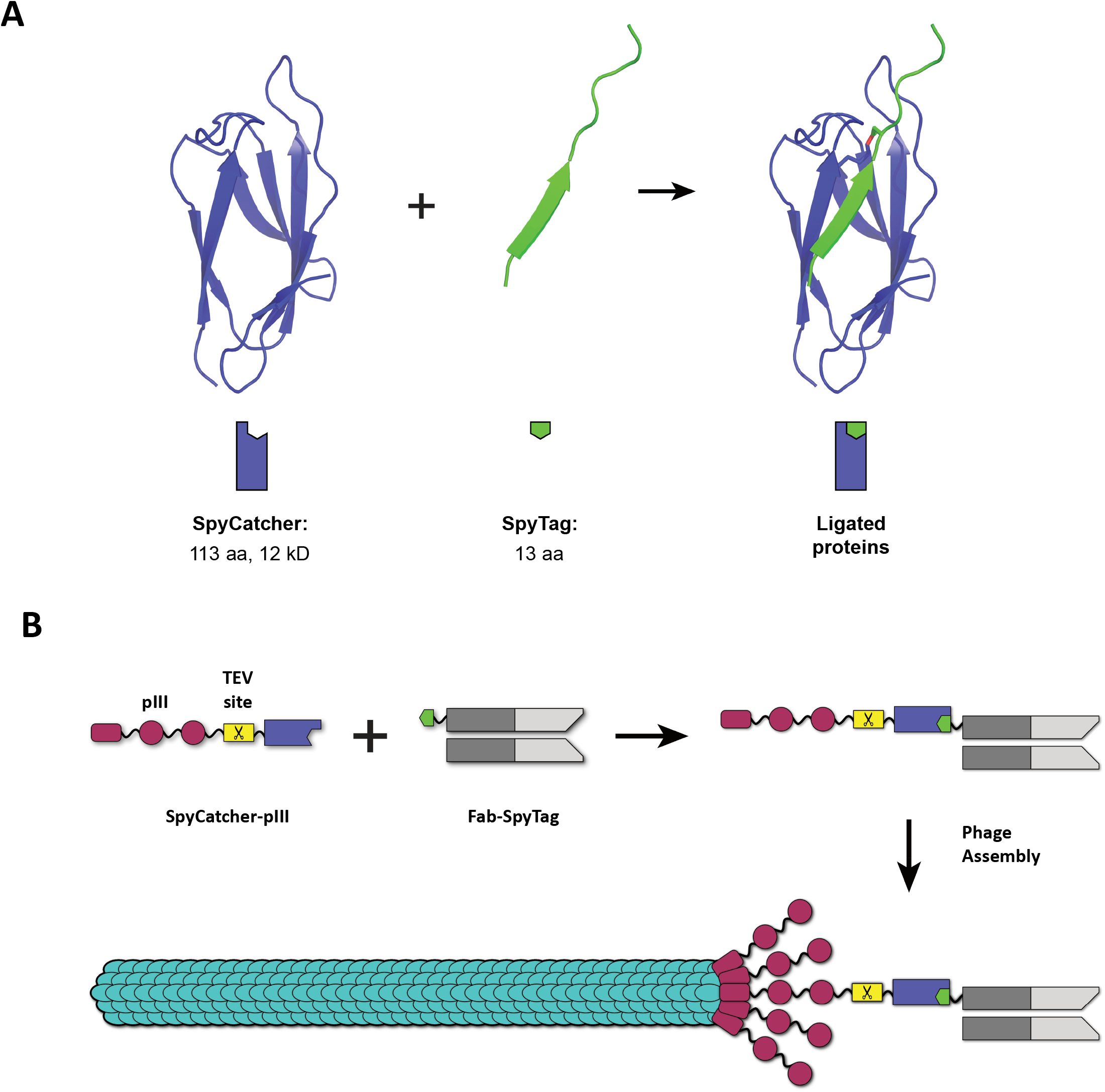
SpyTag/SpyCatcher and the concept of SpyDisplay. **A:** SpyCatcher (violet) and SpyTag (green) react spontaneously to form an isopeptide bond (red). Structures from PDB 4MLI (Li et al., 2014). **B:** In SpyDisplay of antibodies, SpyCatcher-pIII reacts with SpyTagged Fab antibody fragments in the periplasm before (or concomitant with) phage assembly, resulting in phages displaying antibodies (shown here for monovalent display).

In this work, we establish a phage display method based on SpyTag/SpyCatcher technology that we refer to as SpyDisplay. We show that SpyTagged Fab fragments expressed in *E. coli* react *in vivo* with coexpressed SpyCatcher-pIII, resulting in Fab-SpyCatcher-pIII fusion proteins that are incorporated into phage particles, thus enabling phage display (Fig. 1B). In SpyDisplay, the library and display system are separated by using a phagemid-based expression vector which encodes the SpyTagged Fab, and a separate genomically integrated and inducible gene encoding SpyCatcher-pIII. This design avoids any subcloning steps for the expression of free Fab and allows the use of smaller phagemids without the pIII gene. We demonstrate the selection of high-affinity Fab fragments from an antibody library using SpyDisplay. Furthermore, we show that SpyDisplay is a straightforward way for N-terminal phage display (where the protein of interest is displayed with a free C-terminus) and is also compatible with the display of cytoplasmic folding proteins, exported to bacterial periplasm via the twin arginine translocase (TAT) pathway.

## Results

### Generation of a SpyCatcher-pIII expressing *E. coli* strain and production of SpyCatcher-displaying phages

To present antibodies on the phage surface via SpyDisplay, SpyCatcher needs to be fused to M13 minor coat protein pIII. We decided to use the improved version SpyCatcher2/SpyTag2 (Keeble et al., 2017), from now on referred to as SpyCatcher/SpyTag. A TEV protease cleavage site was placed between SpyCatcher and pIII to allow mild and affinity-independent elution of bound phages by proteolytic cleavage (Ward et al., 1996). The SpyCatcher-TEV-pIII expression cassette was integrated into the *E. coli* genome to avoid using an additional plasmid. The SpyCatcher-pIII gene under control of the arabinose promotor was integrated at the *araBAD* locus of TG1 *E. coli* (Fig. 2A), replacing the endogenous *araBAD* genes required for arabinose metabolism (Guzman et al., 1995). In this new bacterial strain, termed SK25, arabinose could be used to induce the expression of SpyCatcher-pIII, as assessed by western blotting (Fig. 2B)

**Figure 2:**
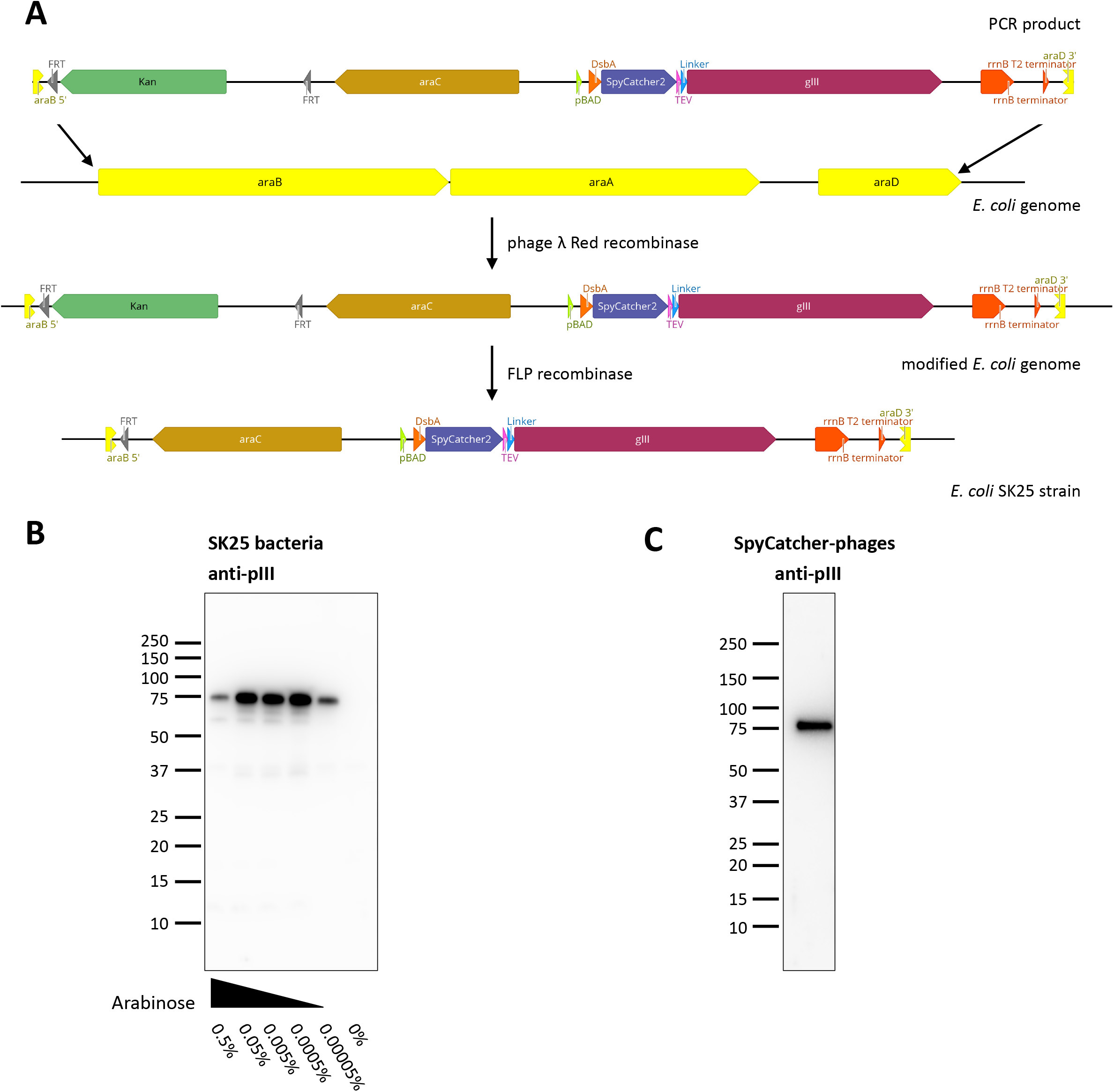
Creation of SpyCatcher-pIII expressing *E. coli* strain SK25. **A:** Phage λ Red recombinase A was used to replace the *araBAD* genes of TG1 *E. coli* with a cassette consisting of a kanamycin resistance gene, *araC*, and SpyCatcher-pIII under control of the pBAD promotor. In the second step, the kanamycin resistance cassette was excised via FLP recombinase. **B**: Expression of SpyCatcher-pIII in SK25 bacteria analyzed by immunoblotting. SK25 cells were grown overnight at 22°C in presence of varying concentrations of arabinose and equal numbers of bacteria were lysed. Lysates were probed with anti-M13-pIII followed by sheep anti-mouse IgG (H/L):HRP. For SpyCatcher-pIII, the apparent molecular weight by SDS PAGE of 75 kDa is higher than the calculated molecular weight of 57 kDa. **C**: Immunoblot of phages produced by infecting SK25 bacteria with Hyperphage. PEG-precipitated phages were separated electrophoretically, immunoblotted, and detection was performed as described in B.

To test the incorporation of SpyCatcher-pIII into the phage capsid, we produced SpyCatcher phages by infecting SK25 cells with Hyperphage (Rondot et al., 2001), a helper phage lacking pIII. Western blotting of these phages with anti-pIII antibody yielded a single, strong band for SpyCatcher-pIII, which migrated at a slightly higher molecular weight than calculated. (Fig. 2C).

### Production of Fab-phages in *E. coli* SK25

To establish antibody phage display via SpyTag/SpyCatcher, Fab fragments of the therapeutic antibodies adalimumab and trastuzumab were cloned into the expression vector pBBx2-F-Spy2-H (Hentrich et al., 2021), which contains an f1 phage replication origin and can therefore be employed as a phagemid. In this phagemid, Fab expression is under control of the *lac* promotor and the heavy chain of the Fab is C-terminally fused to three consecutive peptide tags: FLAG-tag, SpyTag2, and His_6_-tag. The phagemids were transformed into *E. coli* SK25 cells and the cells were subsequently infected with VCSM13 helper phage. Monovalent Fab-phages displaying not more than one Fab antibody fragment per phage particle were produced using arabinose to induce expression of SpyCatcher-pIII and IPTG to induce expression of Fab. In parallel, polyvalent Fab-phages were produced using Hyperphage and analogous induction of expression with arabinose and IPTG. Phages were purified and concentrated by polyethylene glycol (PEG) precipitation from the supernatants of overnight cultures. Initially, phage titers of mono- and polyvalent Fab-phages measured by spot titration differed by three orders of magnitude and were 3.1 × 10^13^ cfu/mL (SD=0.5 × 10^13^ cfu/mL) for monovalent Fab-phages and 6.5 × 10^9^ cfu/mL (SD=1.9 × 10^9^ cfu/mL) for the polyvalent ones. It has been described that fusions to the N-terminus of pIII decrease phage infectivity (Loset et al., 2008) and we suspected this to be the explanation for the apparent low titer of the polyvalent Fab-phages. Therefore, to allow better comparison of titers, Fab-phage particles were digested with TEV protease prior to spot titration, cleaving the Fab-SpyCatcher fusion off the pIII protein. After treatment with TEV protease, the titer of the monovalent phages was unchanged 2.8 × 10^13^ cfu/mL (SD=0.8 × 10^13^ cfu/mL) while the polyvalent titer was 100-fold higher (7.2 × 10^11^ cfu/mL; SD=1.5 × 10^11^ cfu/mL) than before protease treatment. The remaining difference of two orders of magnitude between regular helper phage and Hyperphage titer has been observed before (Loset et al., 2008) and is likely caused by steric hindrance of the modified pIII proteins during phage assembly. Western blots of monovalent and polyvalent SpyDisplay Fab-phages confirmed that Fab-SpyCatcher-pIII fusions were successfully inserted into the phage particles (Fig 3A). As wildtype pIII is incorporated faster into the phage coat than modified pIII, monovalent phages had a much lower ratio of Fab-pIII to wildtype pIII (about 4%, estimated by densitometry) than the polyvalent phages (Fig 3A). Since an estimated 4% of all pIII proteins carry a Fab and there are 5 copies of pIII per phage particle, approximately 20% of all phages display Fab on their surface in the monovalent setup, a range similar to that of other monovalent phage display libraries (Rothe et al., 2008). As expected, polyvalent Fab-phages generated in absence of wildtype pIII exhibited a much higher Fab display rate. However, even in polyvalent display, only roughly 80% of the SpyCatcher-pIII was coupled to a full length Fab, whereas the remaining SpyCatcher-pIII was coupled to what we presume to be a short degradation product of the Fab, caused by cleavage within the CH1 domain of the Fab heavy chain (Robinson et al., 2015). We further evaluated the phages by ELISA with anti-pVIII-HRP as detection reagent. By coating cognate and irrelevant antigens, we confirmed that the displayed Fabs were functional and specific (Fig. 3B). To test the elution of SpyDisplay phages by proteolysis, phages bound to immobilized antigen were incubated with TEV protease or control buffer for 30 minutes, and residual bound phages after washing were quantified by ELISA, confirming the effectiveness of our elution protocol (Fig. 3C, Suppl. Fig. S1).

**Figure 3:**
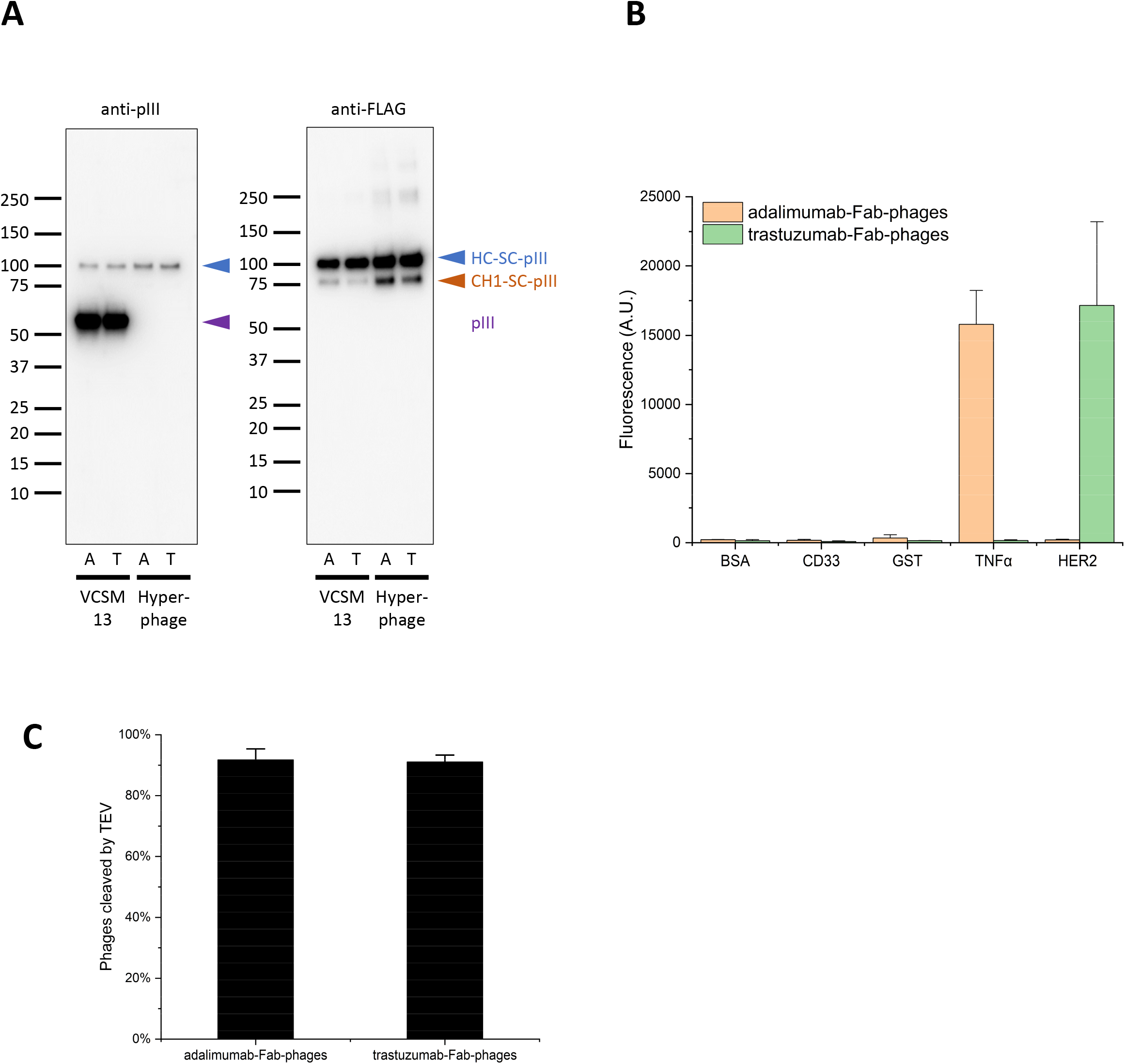
Production of Fab-phages. **A:** Immunoblots of SpyDisplay phages displaying Fabs of adalimumab (A) or trastuzumab (T) and produced with VCSM13 or Hyperphage as helper phage. Detection was performed with anti-M13-pIII followed by sheep anti-mouse IgG (H/L):HRP or anti-FLAG-HRP. Bands corresponding to the heavy chain fusion, degraded heavy chain fusion, and wildtype pIII are marked. **B:** ELISA with polyvalent Fab-phages on cognate and irrelevant antigens, detection with anti-pVIII-HRP. **C:** Fractions of monovalent Fab-phages eluted from MaxiSorp plates after treatment with TEV Protease for 30 minutes, relative to buffer control.

### SpyTag/SpyCatcher-based Fab-phage assembly occurs intracellularly

For coupling of phenotype to genotype, it is essential that the ligation of SpyCatcher-pIII to the SpyTagged Fab takes place inside the bacterial cell. If phage particles presenting free SpyCatcher-pIII were secreted into the supernatant, ligation with Fabs released from other clones in the Fab-phage production culture could occur. This would lead to phages presenting antibodies different from the ones encoded by their phagemid. These scenarios can be tested by spiking experiments, in which SK25 bacteria expressing an antibody *A* are mixed in large excess with bacteria expressing an antibody *B*, and Fab-phages are produced in the mixed culture. For example, at 100:1 mixing ratio of *A* and *B*, and assuming equal Fab and phage production rates, the ratio of phagemids encoding antibodies *A* and *B* in the phage population is expected to be *A*:*B* = 100:1, and the ratio of free antibody released to the medium during phage production would also be *A*:*B* = 100:1.

If coupling occurred exclusively intracellularly, 100% of the phages displaying antibody *B* would carry the correct genotype *B*. On the other end of the spectrum, if coupling occurred exclusively in the medium, one would expect the following distribution: 98.01% of phages displaying *A* and encoding *A* (*A-A*), 0.99% displaying *B* and encoding *A* (*B-A*), 0.99% displaying *A* and encoding *B* (*A-B*), 0.01% displaying *B* and encoding *B* (*B-B*). Therefore, 99% of phages displaying antibody *B* (*B-A* and *B-B*) would carry the wrong genotype *A* (0.99% : 0.01% of total phages). Identifying the genotype of phages displaying the underrepresented antibody *B* is thus a sensitive method to assess the degree of extracellular coupling of Fab to phage.

We therefore mixed exponentially growing cultures of SK25 cells expressing Fabs of therapeutic antibodies adalimumab and trastuzumab in ratios of either 100:1 or 1:100 immediately before superinfection with VCSM13 helper phage. Fab-phages were produced overnight and used for one round of panning on the cognate antigen of the underrepresented antibody. As control for unspecific binding, pure adalimumab and trastuzumab Fab-phages were produced separately, quenched with SpyTag3 peptide to saturate potentially free SpyCatcher sites, and also mixed in volume ratios of 100:1 and 1:100. Phages eluted after the panning round were rescued in SK25 cells and plated. 95 clones were picked from each condition and sequenced. Selection of trastuzumab phages on ErbB2 yielded 100% correct clones (95/95) while selection of adalimumab-phages on TNFα resulted in 96% correct clones (91/95). In the corresponding controls, 99% and 100% correct clones were found, respectively (94/95; 95/95).

The results of both variations of the experiment are in strong agreement with a vast preponderance of intracellular ligation of Fabs to SpyCatcher-pIII. The small number of incorrect genotypes found is similar to the control and is therefore most likely due to unspecific phage binding not unusual in phage display (Miersch et al., 2015).

### SpyDisplay enables N-terminal display and display mediated via the TAT pathway

To display a protein N-terminally on phages, i.e. with a free and unmodified C-terminus, it should be sufficient to attach the SpyTag to the N-terminus of the displayed protein, as SpyTag-SpyCatcher ligation is agnostic towards the position of the tag (Zhang et al., 2013). To test the efficiency of N-terminal SpyDisplay, we added SpyTag to the N-terminus of maltose binding protein (MBP) and of a well characterized anti-fluorescein single chain (scFv) antibody (Honegger et al., 2005). Western blotting of phages produced with Hyperphage showed good display rates, with about 95% of all pIII proteins carrying the displayed protein (Fig 4A, left). Furthermore, we confirmed correct folding of the scFv antibody by testing its capacity to bind its antigen via ELISA (Fig. 4A, right). These data confirm that N-terminal display with SpyDisplay is possible and indeed well-working.

**Figure 4:**
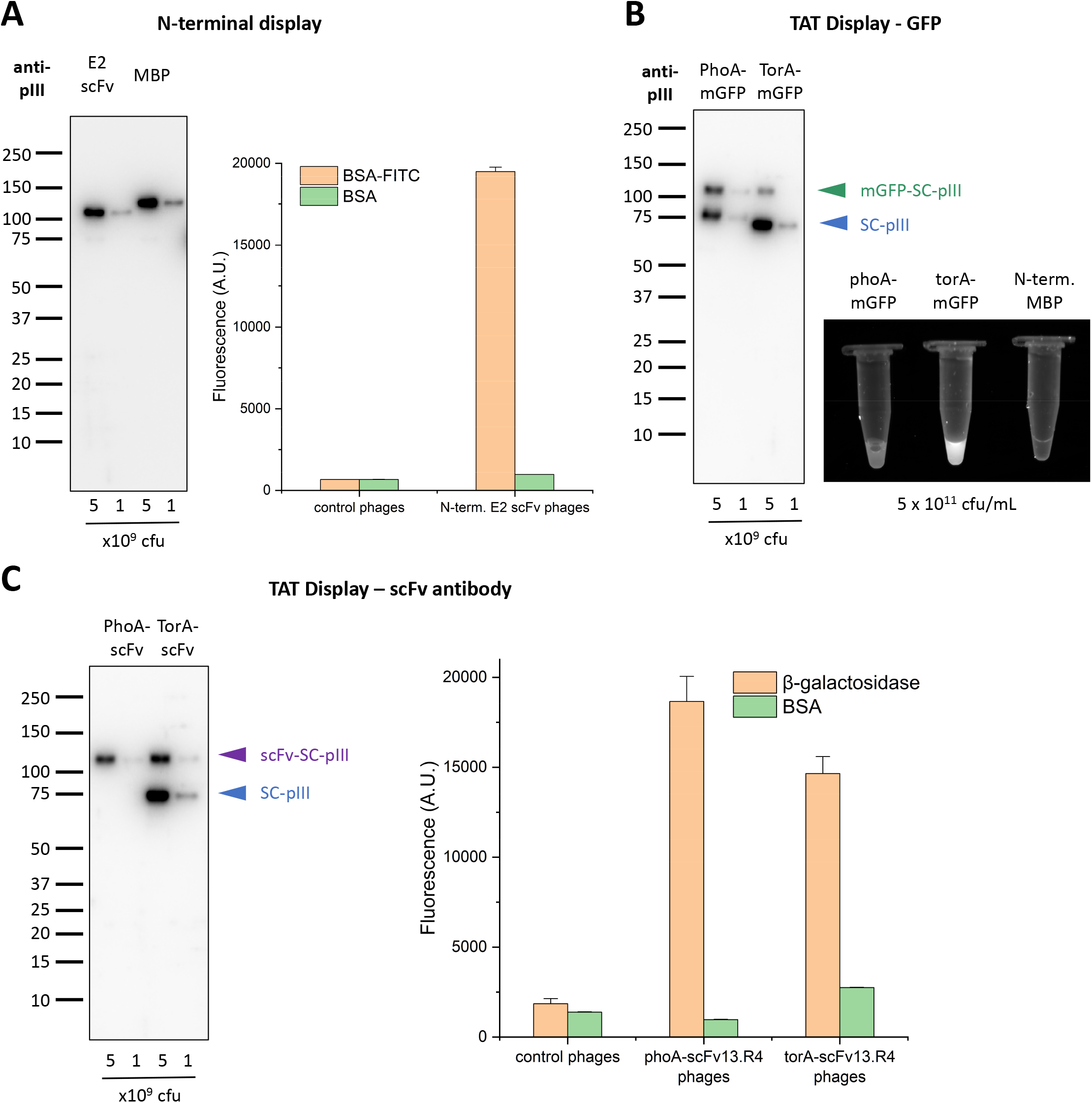
Versatility of display setups with SpyDisplay. **A**: Left: Immunoblot analysis of polyvalent SpyDisplay phages displaying the E2 scFv or MBP with an N-terminal SpyTag. Detection was performed with anti-M13-pIII followed by sheep anti-mouse IgG (H/L):HRP. Right: ELISA of polyvalent phages displaying N-terminally SpyTagged E2 scFv on control (BSA) or cognate antigen (BSA-FITC), in comparison to control phages (displaying N-terminally SpyTagged MBP). **B**: Left: Immunoblot analysis of polyvalent SpyDisplay phages displaying mGFP-SpyTag with a PhoA or TorA leader peptide. Detection performed with anti-M13-pIII followed by sheep anti-mouse IgG (H/L):HRP. Right: Fluorescence image of GFP-phages and control phages at equal concentrations (5 × 10^11^ cfu/mL) in microcentrifuge tubes. **C**: Left: Immunoblot analysis of polyvalent SpyDisplay phages displaying scFv13.R4-SpyTag with a PhoA or TorA leader peptide. Detection performed with anti-M13-pIII followed by sheep anti-mouse IgG (H/L):HRP. Right: ELISA of polyvalent phages displaying scFv13.R4 with on control (BSA) or cognate antigen (β-galactosidase), in comparison to control phages.

Another benefit of SpyDisplay could be the possibility to display proteins that fold in the cytoplasm. In addition to the Sec-dependent protein translocation pathways, where the proteins fold in the periplasm, bacteria possess the twin-arginine translocase (TAT) pathway (Palmer and Berks, 2012). In this pathway, proteins fold in the cytoplasm and are transported in folded state across the plasma membrane into the periplasm. However, using the TAT pathway to display proteins fused to full-length pIII has been challenging in the past, as periplasmic export of full-length pIII is incompatible with this translocation pathway, possibly due to multiple internal disulfide bonds (Speck et al., 2011). We expressed two proteins with a TAT pathway signal sequence, monomeric GFP (mGFP) and an antibody that can fold both in the cytoplasm or periplasm (scFv13.R4) against β-galactosidase (Fisher et al., 2008). GFP has been shown to only efficiently mature its fluorophore when expressed in the cytoplasm but not in the periplasm (Thomas et al., 2001). Indeed, when mGFP-SpyTag equipped with a TorA TAT signal sequence was expressed in SK25 cells superinfected with Hyperphage, strongly fluorescent phages were produced, whereas, at the same phage particle concentration, control PhoA-mGFP-SpyTag (Sec pathway) phages showed only weak fluorescence (Fig. 4B, right), demonstrating the efficacy of TAT pathway-driven SpyDisplay. When analogously expressed in SK25 superinfected with Hyperphage, intracellularly folding TorA-scFv13.R4-SpyTag showed similar activity in ELISA as periplasmatically folding PhoA-scFv13.R4-SpyTag (Fig. 4C). Western blot analysis (Figs 4B, 4C) revealed relatively modest display rates via the TAT pathway (about 10% for mGFP and 30% for the scFv in relation to total pIII). This is probably caused by the TAT pathway not exporting sufficient protein into the periplasm to fully occupy all SpyCatcher sites on the polyvalent phages. The apparent free SpyCatcher sites in the case of PhoA-mGFP display are in fact coupled to a small peptide containing FLAG and SpyTag, caused by degradation of mGFP in the periplasm. This degradation does not occur when mGFP is displayed using the TAT pathway (Suppl. Fig. S2 and slight band shift in Fig. 4B). The presence of excess free SpyCatcher with TAT-based display suggests that the TAT pathway may have lower efficiency than Sec-based export, at least during phage production. Therefore, monovalent display might be best suited for TAT-based display.

### Selection of antibodies from a SpyDisplay Fab-phage library

To test the efficiency of SpyDisplay for selecting high-affinity antibodies, we used a subset of our in-house constructed human Fab library termed Pioneer. The antibodies in this library were of the germlines IGHV1-69 and IGLV3-1, with all 6 complementarity determining regions (CDRs) diversified. This Fab library was cloned in the pBBx2-F-Spy2-H expression vector and was transformed into SK25 cells, yielding 1.3 × 10^11^ transformants. Monovalent Fab-phages were produced by infection of library transformants with VCSM13.

We performed SpyDisplay pannings against mGFP and against the paratope of the therapeutic antibody sarilumab to generate anti-idiotype antibodies. Monovalent display was used for all panning rounds with the aim of selecting high-affinity antibodies. Both antigens were immobilized on MaxiSorp plates in two different ways, by passive adsorption and by using biotinylated antigens on preimmobilized streptavidin/neutravidin, thus yielding 4 independent panning setups. After three rounds of panning, the panning output was transformed into *E. coli* SK13, a bacterial strain optimized for the expression of SpyTagged Fabs by removal of two proteases which cleave SpyTag2 in the periplasm (Hentrich et al., 2021).

To screen for antigen binding clones, 368 randomly picked single colonies from each panning (i.e. 736 colonies per antigen) were grown in 384-well plates, and Fab antibodies were expressed overnight. The next day, bacteria were lysed, and antibody-containing lysates were tested by ELISA. 472 clones (64.1%) from the mGFP pannings and 550 clones (74.7%) from the sarilumab pannings showed ELISA signals of at least tenfold over background (Fig 5A). For passively adsorbed antigens, 30 clones with the highest ELISA signal were sequenced for each target. From pannings on biotinylated antigens, 95 clones with the highest ELISA signals were further screened for a low k_off_-rate via bio-layer interferometry (BLI) (Ylera et al., 2013), and for each antigen the 20 clones with the lowest k_off_-rate were sequenced. For mGFP, sequencing a total of 50 clones resulted in 34 unique antibodies (68%), whereas for sarilumab, 23 unique antibodies (46%) were found. Next, all unique Fabs were expressed in 50 mL cultures, purified (Knappik and Brundiers, 2009), and their monovalent affinities were measured by BLI on immobilized antigen. As mGFP-coated sensors could not be regenerated, anti-mGFP antibodies were first measured at a single concentration, and only for the top 20 antibodies a full kinetic measurement was performed. 98 % of all antibodies measured (20 anti-mGFP and 23 anti-sarilumab) had affinities lower than 10 nM, and 37% had affinities lower than 1 nM. The best antibodies against mGFP and sarilumab had affinities of 40 pM and 24 pM, respectively (Fig 5B, Supp. Fig S3: Sensorgrams of all antibodies found). The measured affinities demonstrate the suitability of SpyDisplay for selection of high-affinity antibodies.

**Figure 5:**
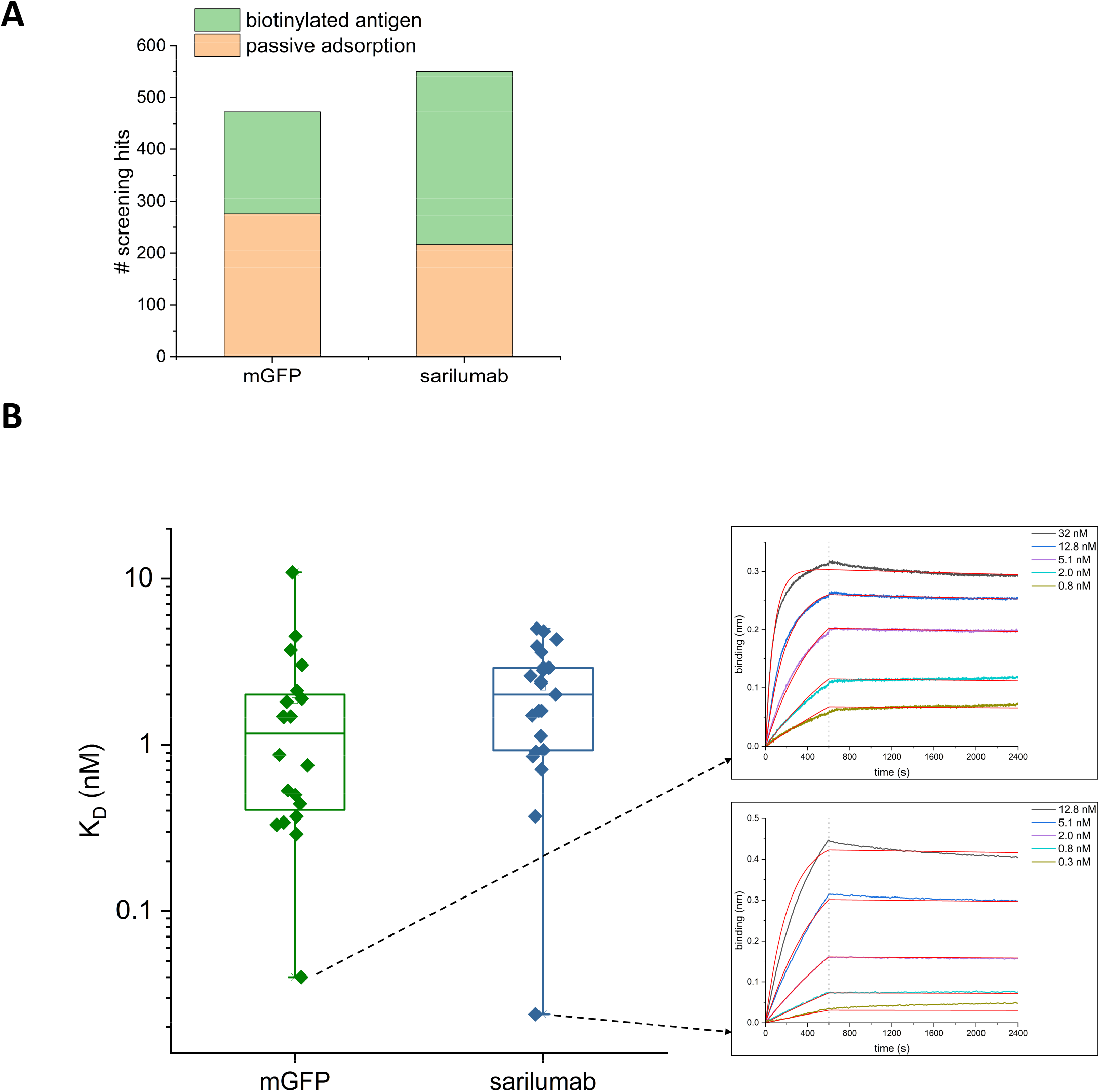
SpyDisplay selections of antibodies from Fab library. **A:** Number of positive hits (>10-fold over background in ELISA on cognate antigen) from screening Fabs generated against mGFP or the paratope of sarilumab (out of 736 tested antibodies per antigen). **B**: Left: Distribution of K_D_ values of sequenced and purified antibodies against mGFP and sarilumab. Right: BLI sensorgrams of the best antibodies against each antigen.

## Discussion

In this study, we have established SpyDisplay as a new phage display method with some important differences to traditional phage display. A key distinction between SpyDisplay and traditional antibody phage display is the separate folding of antibody and coat protein, which may be beneficial for the display of correctly folded antibodies. In practical terms, the use of the library phagemid as expression plasmid saves time by eliminating a molecular cloning step before or after screening, which is laborious, especially if many pannings are performed in parallel. Alternative methods for avoiding subcloning such as using an amber stop codon between Fab and pIII have been shown to deliver sufficient free Fab for screening and small-scale expressions. However, due to low expression yields, production of larger amounts of Fab still necessitates the use of a dedicated expression vector (Chasteen et al., 2006). Having a phagemid without pIII has a further advantage during library construction: as transformation efficiency inversely correlates with plasmid size, larger libraries can be generated by using the smaller SpyDisplay phagemid which lacks the pIII gene (1230 bp) (Hanahan, 1983). Additionally, our optimized SpyDisplay protocol allows to perform one panning round in a single day without the need for plating the output, which is especially important for high throughput antibody selections. Finally, SpyDisplay enables selection of antibodies with high affinity.

Monovalent and polyvalent phage display are commonly implemented by switching the type of helper phage. Monovalent display is crucial for the selection of high-affinity antibodies, whereas polyvalent display provides avidity. The efficiency of monovalent SpyDisplay for the selection of high-affinity antibodies is demonstrated by the selection of antibodies in the low pM range in our test pannings. Polyvalent SpyDisplay has been implemented by using Hyperphage and indeed showed high display rates. We envision that, in the context of Fab library panning, polyvalent SpyDisplay will facilitate the selection of antibodies against challenging targets like carbohydrates (Mazor et al., 2010). Affinities of such antibodies can later be improved by an optional affinity maturation process.

We have shown that the displayed antibody and the anchoring SpyCatcher-pIII protein, which are translated independently of each other, form a fusion protein exclusively in the periplasm. Optimized periplasmic secretion signal peptides for each polypeptide chain can be used. In this implementation of antibody SpyDisplay, three different signal peptides are used: OmpA and PhoA for light and heavy antibody chains, and DsbA for the SpyCatcher-pIII fusion. Furthermore, we showed that the Twin Arginine Translocation (TAT) pathway is compatible with SpyDisplay, enabling the cytoplasmic folding of the displayed proteins. The display rate via the TAT pathway could potentially be increased through optimization of the TAT signal peptide. We anticipate that TAT-based display will be useful for selecting intrabodies, antibodies that can fold in the cytoplasm, without the need for specialized intrabody libraries.

Mazor et al. also achieved a separation of the expression of antibody and coat protein by forming a complex between an IgG and Fc-binding ZZ-domain-pIII fusion (Mazor et al., 2010). Similarly, the Jun/Fos leucine zipper has been used to link the displayed protein to pIII (Paschke and Hohne, 2005). Since these methods are non-covalent, the complex might dissociate during stringent washing steps required for the selection of high-affinity antibodies. However, it is possible to stabilize the leucine zipper by introducing a disulfide bond between the dimerized proteins (Strauch and Georgiou, 2009). CysDisplay is another phage display method to express the antibody separately from pIII and relies on the spontaneous formation of disulfide bonds between engineered free cysteines at the C-terminus of Fab heavy chain and the N-terminus of pIII. While this method has proven powerful and became the basis of the HuCAL platform (Rothe et al., 2008), significant side reactions in this setup are the formation of pIII homodimers as well as Fab homodimers, making the control of the display rate much less straightforward compared to the specific SpyTag-SpyCatcher reaction.

In a previous study (Hentrich et al., 2021), we showed that efficient periplasmic expression of SpyTagged Fabs is only possible in a protease knockout strain of *E. coli*, termed SK13. However, the proteases responsible for SpyTag cleavage do not seem to harm the assembly of Fab-SpyCatcher-pIII-phages. This suggests that the ligation of SpyTagged Fab to SpyCatcher-pIII occurs sufficiently fast *in vivo* and that SpyTag is protected from cleavage by becoming part of the structured Ig-like fold of the reconstituted SpyTag-SpyCatcher fusion. This is consistent with the greatly enhanced thermal stability of SpyCatcher after coupling to SpyTag (Hentrich et al., 2021). Therefore, the SK25 (an F’ strain without protease knock-outs) can be used for all SpyDisplay panning steps. For screening and purification, the selected phagemids are transformed into SK13 to express high concentrations of soluble SpyTagged Fabs. SK13 is an F-strain and thus also avoids unintended infection with contaminating phages, which could lead to the expression of other antibodies in parallel. The resulting Fabs are fully compatible with SpyTag-based modular antibody assembly, enabling rapid site-specific labeling and change of oligomeric state using prefabricated SpyCatcher modules (Hentrich et al., 2021). This enables screening in mono-and bivalent format in parallel and greatly accelerates further antibody characterization, thus providing a substantial economic benefit.

We have established SpyDisplay here mainly for the selection of antibodies, however we do not expect any limitations regarding the displayable proteins, provided they contain a SpyTag and can be transported to the periplasm. The SpyTag of the displayed protein can be inserted at any accessible position within a protein (Zakeri et al., 2012), providing manifold options for the topology of protein display on the phage. In particular, SpyTag fused N-terminally to the displayed protein enables selection of proteins which require an unmodified C-terminus, which is not possible with the standard pIII-fusion approach. SpyTag/SpyCatcher seems furthermore well suited to display proteins in other selection systems. For example, both yeast display (Kajiwara et al., 2021) and bacterial display (Gallus et al., 2022; Gallus et al., 2020) via SpyTag/SpyCatcher have been demonstrated. The physical separation of the SpyTagged library from the other elements of the display system as shown here enables switching between display methods, provided the vector stays compatible with the host expression system. By utilizing a dual expression vector compatible with both *E. coli* and yeast expression (Patel et al., 2011), it would be possible to seamlessly switch from phage to yeast display simply by transforming the output of phage display into a SpyDisplay-compatible yeast strain displaying SpyCatcher. Similarly, a compatible plasmid would allow for a transition from phage display to mammalian display (Tesar and Hotzel, 2013). Such an approach could combine the benefits of these popular protein selection systems, namely the large library size of phage display and the ability to perform eukaryotic expression and flow cytometry sorting of the selection output in yeast or mammalian cells.

Other protein ligation technologies besides SpyTag/SpyCatcher have been established and could be used to establish analogous ‘covalent capture’ display methods. Such technologies include SnoopTag/SnoopCatcher (Veggiani et al., 2016), DogTag/DogCatcher (Keeble et al., 2022), SilkTag/SilkCatcher (Fan et al., 2022), split inteins (Vila-Perello and Muir, 2010), or enzyme-based ligation technologies such as ones based on sortase (Schmohl and Schwarzer, 2014), butelase (Nguyen et al., 2014), or peptiligase (Toplak et al., 2016).

*In vitro* selections are often hindered by biases such as the accumulation of truncated sequences or selection not based on affinity (Plessers et al., 2021). The high screening hit rates, high diversities, and subnanomolar affinities of the antibodies selected by SpyDisplay suggest that SpyDisplay is resilient against such biases. This study demonstrates that SpyDisplay has the potential to significantly improve selection campaigns.

## Materials

### Plasmids

pBBx2-F-Spy2-H (Hentrich et al., 2021),, pKD46 (Datsenko and Wanner, 2000), pCP20 (Datsenko and Wanner, 2000), pKD13 (Datsenko and Wanner, 2000), pET28a_SpyCatcher2 (Hentrich et al., 2021), pACYC177 (Chang and Cohen, 1978)

### Oligonucleotides

161_SKE, 175_SKE, 185_SKE, 186_SKE, 164_SKE, 165_SKE, 178_SKE, 179_SKE, 180_SKE, 181_SKE, 182_SKE, 183_SKE (all this study, nucleotide sequences shown in the supplementary information)

### Strains and phages

*E. coli* TG1, *E. coli* SK25 (this study), *E. coli* SK13 (Hentrich et al., 2021)

VCSM13 (Agilent), Hyperphage (Progen)

### Antibodies

Anti-M13-pIII monoclonal antibody (E8033S, New England Biolabs), sheep anti-mouse IgG (H/L):HRP (AAC10P, Bio-Rad), anti-FLAG-Tag (M2) IgG-HRP (A8592, Sigma), anti-M13 bacteriophage coat protein g8p (ab9225, Abcam), goat anti-human IgG F(ab’)2 (STAR126, Bio-Rad), anti-M13-pVIII-HRP (Cytiva 27-9421-01, Sigma)

## Methods

### *E. coli* cultivation

*E. coli* strains were cultivated in 2xYT medium supplemented with glucose (Glc, varying concentrations), arabinose (Ara, varying concentrations), isopropyl β-D-1-thiogalactopyranoside (IPTG, 0.25 mM), kanamycin (Kan, 50 µg/mL), and/or chloramphenicol (Cam, 34 µg/mL). Cells were grown on orbital shakers in Erlenmeyer flasks at 250 rpm (Fab-phage production) or in 24 deep-well blocks at 400 rpm (panning) at temperatures between 22°C and 37°C. For generation of SK25, plates were incubated at 42°C for plasmid curing. For selection of the F plasmid in TG1 derivatives, cells were plated on M9 agar.

### Plasmid cloning

For selection and soluble expression of selected binders from SpyDisplay pannings, pBBx2-F-Spy2-H (Hentrich et al., 2021) was used.

### Human antibody phage display library

A subset of the Pioneer library (Bio-Rad) was used for phage display, which contained human Fab antibodies with the heavy chain germline IGHV1-69 and the light chain germline IGLV3-1. The CDR diversity was generated by gene synthesis and CDRs were cloned into the plasmid pBBx2-F-Spy2-H in successive steps.

### Generation of *E. coli* SK25

The method used for generation of *E. coli* SK25 is based on the λ-Red-recombinase system described by Datsenko and Wanner (Datsenko and Wanner, 2000). First, the PCR-product (sequence in supplementary information) consisting of the SpyCatcher2-pIII expression cassette with the FRT-Kan-FRT cassette and *araC* was amplified with oligonucleotides 161_SKE and 175_SKE using the template for SK25 generation. PCR-products of correct size were gel-purified using Wizard SV Gel and PCR Clean-Up Kit (Promega). *E. coli* TG1 cells harboring the helper plasmid pKD46 were made electrocompetent (in presence of 50 mM arabinose instead of 1 mM arabinose) and transformed with the PCR product as described (Datsenko and Wanner, 2000). Mutants were identified by colony PCR using the oligos 185_SKE and 186_SKE. After helper plasmid curing, the KanR mutant was transformed with pCP20 for excision of the KanR cassette via Flp-FRT recombination. After curing of pCP20, correct excision of the KanR cassette was analyzed by colony PCR using primers 164_SKE and 165_SKE. Final verification of the correct integration of the SpyCatcher2-pIII expression cassette was performed by sequencing of the final colony PCR product with primers 164_SKE, 165_SKE, 178_SKE, 179_SKE, 180_SKE, 181_SKE, 182_SKE, and 183_SKE. Maintenance of the F plasmid was controlled after each mutagenesis step by plating on M9 agar.

### Colony PCR

Fresh colonies were picked and directly resuspended in 16 µL of the PCR mastermix. PCR was performed with hot start *Taq* DNA polymerase (New England Biolabs) according to the manufacturer’s instructions. For generation of *E. coli* SK25, wildtype *E. coli* TG1 was always tested side-by-side as control.

### Fab-phage production

2xYT/1%Glc/Cam medium was inoculated from a glycerol stock of *E. coli* SK25 harboring the antibody sublibrary to an OD_600_ of 0.1 and cultivated at 37°C and 220 rpm in 2 L shake flasks. At OD_600_ of 0.5 to 0.8, helper phages were added (VCSM13 in monovalent SpyDisplay, Hyperphage in the polyvalent setup), followed by incubation at 37°C for 45 min without shaking and another 45 min at 220 rpm. After helper phage infection, medium was changed (2xYT/Kan/Cam/0.25 mM IPTG/0.002% arabinose) for overnight production of Fab-phages at 22°C and 220 rpm. For small scale preparations (< 2 L), phage-containing supernatants were harvested by centrifugation (6,000 x *g*, 30 min, 4°C). After filtration (0.22 µm) phages were precipitated with ¼th volume of 20% (w/v) PEG 6000/2.5 M NaCl for 60 min on ice, followed by centrifugation (13,000 x *g*, 30 min, 4°C). The phage pellet was resuspended in PBS/20% glycerol and stored at −80°C. For large scale preparations (> 2 L), phages were filtered and concentrated using a SartoJet Membrane Pump (Sartorius) with Sartocon Slice Hydrosart cassettes (0.45 µm for cell removal and 100 kDa cutoff for phage concentration; Sartorius), followed by the final precipitation with PEG/NaCl as described above.

### Phage ELISA

96-well plates (Maxisorp F96, Thermo Fisher Scientific) were coated with anti-M13 bacteriophage coat protein g8p antibody or goat anti-human IgG F(ab’)2 in PBS overnight at 4°C. Plates were washed five times with TBST (TBS with 0.05% (v/v) Tween 20) after coating and blocking with 33% (v/v) ChemiBLOCKER (Merck Millipore) in TBST for 1 h at RT. Next, serial dilutions of Fab-phages in 33% (v/v) ChemiBLOCKER/TBST were transferred to the blocked plates and incubated for 1 h at RT. After washing (ten times with TBST), bound Fab-phages were detected with anti-M13-pVIII-HRP diluted 1:5,000 in 33% (v/v) ChemiBLOCKER/TBST (1 h, RT), followed by a final wash (ten times with TBST) and addition of QuantaBlu fluorogenic peroxidase substrate (Thermo Fisher Scientific). Fluorescence (excitation at 320 ± 25 nm, emission at 430 ± 35 nm) was measured with an Infinite 200 microplate reader (Tecan). To measure the efficiency of phage elution by TEV proteolysis, serially diluted phages were bound on cognate antigens, followed by incubation with TEV Buffer (1 µg/mL TEV protease in 50 mM Tris pH 8.0, 0.5 mM EDTA, 1 mM DTT, 0.5% (w/v) BSA) or control buffer (same buffer but without TEV protease). After washing, an ELISA with anti-pVIII-HRP was performed to quantify the remaining bound phages after elution. The linear regions of the digest and corresponding control curves were fitted with a linear equation with shared Y-axis intercept (Suppl Fig S1). To determine the efficiency of cleavage, the slope of the TEV digestion condition was divided by the slope of the corresponding control and fit errors were propagated.

### Immunoblotting

Western blots were performed as described (Hentrich et al., 2021). Briefly, different dilutions of phage preparations were run on SDS-PAGE and afterwards blotted onto PVDF membranes via semi-dry transfer. For detection of pIII and derivatives, blots were incubated with anti-M13-pIII monoclonal antibody, followed by washing and incubation with sheep anti-mouse IgG (H/L):HRP. Alternatively, anti-FLAG-Tag (M2) IgG-HRP was used for detection of FLAG-tagged proteins. Clarity ECL substrate (Bio-Rad) was used as HRP substrate and a Chemidoc MP imager (Bio-Rad) was used for signal detection.

### Spot titration

Titer of phage preparations was determined by spot titration. (Fab)-phage preparations were serially ten-fold diluted in 2xYT-medium in a 96-well microtiter plate (ThermoFisher Scientific) in triplicates. Equal amounts of freshly grown *E. coli* SK25 with an OD_600_ of 0.5 to 0.8 were added and the mixture was incubated for 30 min at 37°C. Afterwards, 5 µL of each dilution was spotted on LB/Cam/Glc agar plates. Titers (colony-forming units (cfu)/mL) were determined after overnight incubation at 37°C.

### Antibody selection with SpyDisplay

Antigens or streptavidin/neutravidin were immobilized overnight on polystyrene plates (MaxiSorp, Thermo Fisher Scientific). Plate blocking was performed with 5% (w/v) milk powder or 5% (w/v) BSA in PBST (PBS with 0.05% (v/v) Tween 20) for 1 hour at RT, followed by incubation with biotinylated antigen where applicable. Blocked Fab-phages were transferred onto the antigen-coated plates and incubated for 3 hours (1st round) or 2 hours (2nd and 3rd round) at RT at 400 rpm. After washing (5 times with TBST), selected Fab-phages were eluted with TEV-buffer (1 µg/mL TEV in 50 mM Tris pH 8.0, 0.5 mM EDTA, 1 mM DTT, 0.5% (w/v) BSA) for 30 min at RT and transferred to 3.5 mL *E. coli* SK25 cells (OD600 = 0.6 – 0.8) in 24-deep-well blocks. To allow phage infection of bacteria, the cultures were incubated at 37°C, 400 rpm for 1 h. Afterwards, chloramphenicol was added to a final concentration of 34 µg/mL, followed by further incubation at 37°C and 400 rpm for 30 min. VCSM13 was added to a final concentration of 10^9^ cfu/mL, followed by incubation at 37°C and 400 rpm for 60 min. Following infection, cells were centrifuged at 2,200 x g at RT for 5 min and resuspended in a larger volume (25 mL Fab-phage expression medium: 2xYT/Cam/Kan/0.25 mM IPTG/0.005% Ara) to ensure the efficacy of the antibiotics (Supp. Fig. S4). Fab-phages were expressed overnight at 30°C and 350 rpm in an orbital shaker (Multitron, Infors HT). The next panning round was performed with Fab-phage-containing supernatants of the overnight cultures. After three rounds of panning, the plasmids of the panning output were isolated via DNA preparation (PureYield Plasmid Miniprep System, Promega) and transformed into chemically competent *E. coli* SK13 (Hentrich et al., 2021) cells.

### Identification of antigen-binding clones via primary screening ELISA

Identification of antigen binding clones was performed as described previously (Jarutat et al., 2006). Briefly, 368 single clones were picked, Fab-fragments were expressed overnight in 384-well microtiter plate (ThermoFisher Scientific), and crude bacterial lysates were used for ELISA screening. 384-well plates (MaxiSorp black, ThermoFisher Scientific) were coated with antigens and controls overnight in PBS at 4°C. Plates were washed five times with PBST after coating and blocking with 5% milk-PBST or 5% BSA-PBST for 1 h at RT. Next, blocked lysates (diluted 1:2 in PBST) were transferred onto the antigen coated plates and incubated for 1 h at RT. After washing (ten times with PBST), bound Fab-fragments were detected with anti-FLAG-tag (M2) IgG-HRP diluted 1:20,000 in 0.5% milk-PBST (1 h, RT), followed by washing (ten times with PBST) and addition of QuantaBlu Fluorogenic Peroxidase Substrate (Thermo Fisher Scientific). Fluorescence (excitation at 320 ± 25 nm, emission at 430 ± 35 nm) was measured with an Infinite 200 microplate reader (Tecan). Antibodies which showed a signal greater than ten-fold over background and did not recognize the negative control proteins were considered as antigen-binding clones. For pannings on passively adsorbed antigen, 30 clones with the highest ELISA signals were sequenced and unique clones were expressed and purified as described below. For pannings on biotinylated antigens, a K_off_ ranking was performed with 95 top hits from primary screening.

### Koff ranking via bio-layer interferometry

For clones derived from pannings on biotinylated antigens, off-rate screening was performed on Octet RED384 and Octet HTX instruments (Sartorius) as described previously (Ylera et al., 2013). Lysates of 95 antibodies showing the strongest signal in primary screening ELISA were tested. Biotinylated antigens were immobilized on streptavidin sensors (SA, Sartorius) with typical immobilization levels of 5 ± 0.5 nm and 1.5 ± 0.5 nm for biotinylated mGFP (lot-specific variations) and 5 ± 0.5 nm for biotinylated sarilumab. Baseline was measured in mock lysates without target-specific antibodies for 5 min, followed by the association phase (7.5 min) in specific lysates. Dissociation rate was again measured in mock lysates (7.5 min). Data were analyzed using a 1:1 interaction model with Octet Analysis Studio software 12.2. Curves with poor fits (R^2^ < 0.96) were excluded from analysis.

### Protein expression and purification

Fab fragments were expressed and purified as described previously (Knappik and Brundiers, 2009). In short, 2xYT/0.1%Glc/Cam medium was inoculated from an overnight culture with *E. coli* SK13 containing the expression plasmids and cultivated at 30°C and 250 rpm until the OD_600_ reached about 0.5, followed by induction with 0.8 mM IPTG and overnight incubation at 27.5°C. Bacterial pellets were lysed with BugBuster (Merck KGaA), supplemented with 20 units/mL Benzonase (Merck KGaA), 2 mg/mL lysozyme (Merck KGaA) and protease inhibitors (Complete EDTA free; Roche) and loaded on Ni-NTA agarose (Qiagen). After washing, Fab fragments were eluted with imidazole-containing buffer (250 mM imidazole, 500 mM NaCl, 20 mM NaH_2_PO_4_ pH 7.4), followed by buffer exchange to PBS via PD10 columns (GE Healthcare). TEV protease was expressed and purified as described (Tropea et al., 2009).

### Affinity determination

Affinities of purified Fab fragments were determined on Octet RED384 and Octet HTX instruments (Sartorius) as described previously (Ylera et al., 2013). Briefly, biotinylated antigens were immobilized on streptavidin sensors (SA, Sartorius). Kinetic parameters were determined with purified Fab fragments at five concentrations ranging from 200 nM to 0.13 nM in running buffer (PBS, 0.1% (w/v) BSA, 0.02% (v/v) Tween 20). The association was measured for 600 s and the dissociation for 300 to 1,800 s, depending on the binding strength. For sarilumab, before each single cycle of the measurement, biosensor surface was regenerated with 10 mM glycine, pH 3.0. In case of mGFP, each Fab dilution was measured on a separate sensor without regeneration. Data were analyzed using a 1:1 interaction model with Octet Analysis Studio software 12.2.

## Supporting information

Supplement sequences and figures

## Abbreviations

CDR: complementarity determining region
cfu: colony-forming unit
Fab: antigen binding fragment of an antibody
FRT: flippase recognition target
HRP: horseradish peroxidase
IgG: immunoglobulin G
IPTG: isopropyl β-D-1-thiogalactopyranoside
κ: kappa light chain of an antibody
λ: lambda light chain of an antibody
MBP: maltose-binding protein
mGFP: monomeric green fluorescent protein
pIII: filamentous phage protein III
TAT: twin arginine translocase
TEV: tobacco etch virus
ELISA: enzyme- linked immunosorbent assay
BLI: biolayer interferometry
PBS: phosphate-buffered saline
TBS: tris- buffered saline
BSA: bovine serum albumin
PEG: polyethylene glycol
scFv: single-chain variable fragment
SD: standard deviation

## Acknowledgements

We thank our colleagues in the Fab production and assay teams for antibody purification and quality control and our colleagues in lab support for production of the library phages. We thank Melissa Wich for critical reading of the manuscript.

## Contributions

S.-J.K., H.H., M.C., C.H., and M.P. performed experiments. S.-J.K., C.H., M.P., and F.Y. designed experiments and analyzed the data. A.K. and F.Y. conceived the project. C.H., S.-J.K., M.P., A.K., and F.Y. wrote the manuscript.

