## Supplement sequences and figures for "SpyDisplay: A Versatile Phage Display Selection System using SpyTag/SpyCatcher Technology"

### Supplementary Information for Kellmann *et al.*, "SpyDisplay: A Versatile Phage Display Selection System using SpyTag/SpyCatcher Technology"

1) Primers used in this study
2) Sequence of PCR cassette for generation of SK25
3) Supplementary figures
4) Supplemental references

#### Primers used in this study

| *Name* | *Sequence* |
|---|---|
| 161_SKE | CGATGGCGATTGCAATTGGCCTCGATTTTGGCAGTGATTCTGTGCGAGCTGTGTAGGCTGGAGCTG CTTC |
| 164_SKE | ACGATGGCGATTGCAATTGG |
| 165_SKE | TCCATCAGATAGCGTTCTGG |
| 175_SKE | TTACTGCCCGTAATATGCCTTCGCGCCATGCTTACGCAGATAGTGTTTATGTCTCATGAGCGGATA CATATTTG |
| 178_SKE | TGACAACTTGACGGCTACATC |
| 179_SKE | GATACCATTCGCGAGCCTCC |
| 180_SKE | TAGCTGGTCTTGTGTTGGCG |
| 181_SKE | TGGGTTCCTATTGGGCTTGC |
| 182_SKE | GCTCTGGTTCCGGTGATTTTG |
| 183_SKE | CAGCCTGATACAGATTAAATCAG |
| 185_SKE | CATCAAGCAGCCAATAATCC |
| 186_SKE | GCTACCCGTGATATTGCTG |

#### Template for amplification of PCR-cassette for generation of SK25

GTGTAGGCTGGAGCTGCTTCGAAGTTCCTATACTTTCTAGAGAATAGGAACTTCGGAATAGGAACTTCAAGATCCCCT
TATTAGAAGAACTCGTCAAGAAGGCGATAGAAGGCGATGCGCTGCGAATCGGGAGCGGCGATACCGTAAAGCACGAGG
AAGCGGTCAGCCCATTCGCCGCCAAGCTCTTCAGCAATATCACGGGTAGCCAACGCTATGTCCTGATAGCGGTCCGCC
ACACCCAGCCGGCCACAGTCGATGAATCCAGAAAAGCGGCCATTTTCCACCATGATATTCGGCAAGCAGGCATCGCCA
TGGGTCACGACGAGATCCTCGCCGTCGGGCATGCGCGCCTTGAGCCTGGCGAACAGTTCGGCTGGCGCGAGCCCCTGA
TGCTCTTCGTCCAGATCATCCTGATCGACAAGACCGGCTTCCATCCGAGTACGTGCTCGCTCGATGCGATGTTTCGCT
TGGTGGTCGAATGGGCAGGTAGCCGGATCAAGCGTATGCAGCCGCCGCATTGCATCAGCCATGATGGATACTTTCTCG
GCAGGAGCAAGGTGAGATGACAGGAGATCCTGCCCCGGCACTTCGCCCAATAGCAGCCAGTCCCTTCCCGCTTCAGTG
ACAACGTCGAGCACAGCTGCGCAAGGAACGCCCGTCGTGGCCAGCCACGATAGCCGCGCTGCCTCGTCCTGCAGTTCA
TTCAGGGCACCGGACAGGTCGGTCTTGACAAAAAGAACCGGGCGCCCCTGCGCTGACAGCCGGAACACGGCGGCATCA
GAGCAGCCGATTGTCTGTTGTGCCCAGTCATAGCCGAATAGCCTCTCCACCCAAGCGGCCGGAGAACCTGCGTGCAAT
CCATCTTGTTCAATCATGCGAAACGATCCTCATCCTGTCTCTTGATCAGATCTTGATCCCCTGCGCCATCAGATCCTT
GGCGGCAAGAAAGCCATCCAGTTTACTTTGCAGGGCTTCCCAACCTTACCAGAGGGCGCCCCAGCTGGCAATTCCGGT
TCGCTTGCTGTCCATAAAACCGCCCAGTCTAGCTATCGCCATGTAAGCCCACTGCAAGCTACCTGCTTTCTCTTTGCG
CTTGCGTTTTCCCTTGTCCAGATAGCCCAGTAGCTGACATTCATCCGGGGTCAGCACCGTTTCTGCGGACTGGCTTTC
TACGTGTTCCGCTTCCTTTAGCAGCCCTTGCGCCCTGAGTGCTTGCGGCAGCGTGAGCTTCAAAAGCGCTCTGAAGTT
CCTATACTTTCTAGAGAATAGGAACTTCGAACTGCAGGTCGACGGATCCCCGGAATGGTACCTGCCTGTCAAATGGAC

GAAGCAGGGATTCTGCAAACCCTATGCTACTCCGTCAAGCCGTCAATTGTCTGATTCGTTACCAATTATGACAACTTG
ACGGCTACATCATTCACTTTTTCTTCACAACCGGCACGGAACTCGCTCGGGCTGGCCCCGGTGCATTTTTTAAATACC
CGCGAGAAATAGAGTTGATCGTCAAAACCAACATTGCGACCGACGGTGGCGATAGGCATCCGGGTGGTGCTCAAAAGC
AGCTTCGCCTGGCTGATACGTTGGTCCTCGCGCCAGCTTAAGACGCTAATCCCTAACTGCTGGCGGAAAAGATGTGAC
AGACGCGACGGCGACAAGCAAACATGCTGTGCGACGCTGGCGATATCAAAATTGCTGTCTGCCAGGTGATCGCTGATG
TACTGACAAGCCTCGCGTACCCGATTATCCATCGGTGGATGGAGCGACTCGTTAATCGCTTCCATGCGCCGCAGTAAC
AATTGCTCAAGCAGATTTATCGCCAGCAGCTCCGAATAGCGCCCTTCCCCTTGCCCGGCGTTAATGATTTGCCCAAAC
AGGTCGCTGAAATGCGGCTGGTGCGCTTCATCCGGGCGAAAGAACCCCGTATTGGCAAATATTGACGGCCAGTTAAGC
CATTCATGCCAGTAGGCGCGCGGACGAAAGTAAACCCACTGGTGATACCATTCGCGAGCCTCCGGATGACGACCGTAG
TGATGAATCTCTCCTGGCGGGAACAGCAAAATATCACCCGGTCGGCAAACAAATTCTCGTCCCTGATTTTTCACCACC
CCCTGACCGCGAATGGTGAGATTGAGAATATAACCTTTCATTCCCAGCGGTCGGTCGATAAAAAAATCGAGATAACCG
TTGGCCTCAATCGGCGTTAAACCCGCCACCAGATGGGCATTAAACGAGTATCCCGGCAGCAGGGGATCATTTTGCGCT
TCAGCCATACTTTTCATACTCCCGCCATTCAGAGAAGAAACCAATTGTCCATATTGCATCAGACATTGCCGTCACTGC
GTCTTTTACTGGCTCTTCTCGCTAACCAAACCGGTAACCCCGCTTATTAAAAGCATTCTGTAACAAAGCGGGACCAAA
GCCATGACAAAAACGCGTAACAAAAGTGTCTATAATCACGGCAGAAAAGTCCACATTGATTATTTGCACGGCGTCACA
CTTTGCTATGCCATAGCATTTTTATCCATAAGATTAGCGGATCCTACCTGACGCTTTTTATCGCAACTCTCTACTGTT
TCTCCATACCCGTCTAGATAACGAGGGCAAAAAATGAAGAAGATCTGGTTAGCGTTAGCTGGTCTTGTGTTGGCGTTT
AGTGCTAGTGCCGCCATGGTAACCACCTTATCAGGTTTATCAGGTGAGCAAGGTCCGTCCGGTGATATGACAACTGAA
GAAGATAGTGCTACCCATATTAAATTCTCAAAACGTGATGAGGACGGCCGTGAGTTAGCTGGTGCAACTATGGAGTTG
CGTGATTCATCTGGTAAAACTATTAGTACATGGATTTCAGATGGACATGTGAAGGATTTCTACCTGTATCCAGGAAAA
TATACATTTGTCGAAACCGCAGCACCAGACGGTTATGAGGTAGCAACTGCTATTACCTTTACAGTTAATGAGCAAGGT
CAGGTTACTGTAAATGGCGAAGCAACTAAAGGTGACGCTCATACTGGATCCAGTGGTAGCGAAAACCTGTACTTTCAA
GGCGGTGGCGGCGGCAGTGGCGGTGGCGGGTCTGCTGAAACTGTTGAAAGTTGTTTAGCAAAACCCCATACAGAAAAT
TCATTTACTAACGTCTGGAAAGACGACAAAACTTTAGATCGTTACGCTAACTATGAGGGCTGTCTGTGGAATGCTACA
GGCGTTGTAGTTTGTACTGGTGACGAAACTCAGTGTTACGGTACATGGGTTCCTATTGGGCTTGCTATCCCTGAAAAT
GAGGGTGGTGGCTCTGAGGGTGGCGGTTCTGAGGGTGGCGGTTCTGAGGGTGGCGGTACTAAACCTCCTGAGTACGGT
GATACACCTATTCCGGGCTATACTTATATCAACCCTCTCGACGGCACTTATCCGCCTGGTACTGAGCAAAACCCCGCT
AATCCTAATCCTTCTCTTGAGGAGTCTCAGCCTCTTAATACTTTCATGTTTCAGAATAATAGGTTCCGAAATAGGCAG
GGGGCATTAACTGTTTATACGGGCACTGTTACTCAAGGCACTGACCCCGTTAAAACTTATTACCAGTACACTCCTGTA
TCATCAAAAGCCATGTATGACGCTTACTGGAACGGTAAATTCAGAGACTGCGCTTTCCATTCTGGCTTTAATGAGGAT
CCATTCGTTTGTGAATATCAAGGCCAATCGTCTGACCTGCCTCAACCTCCTGTCAATGCTGGCGGCGGCTCTGGTGGT
GGTTCTGGTGGCGGCTCTGAGGGTGGTGGCTCTGAGGGTGGCGGTTCTGAGGGTGGCGGCTCTGAGGGAGGCGGTTCC
GGTGGTGGCTCTGGTTCCGGTGATTTTGATTATGAAAAGATGGCAAACGCTAATAAGGGGGCTATGACCGAAAATGCC
GATGAAAACGCGCTACAGTCTGACGCTAAAGGCAAACTTGATTCTGTCGCTACTGATTACGGTGCTGCTATCGATGGT
TTCATTGGTGACGTTTCCGGCCTTGCTAATGGTAATGGTGCTACTGGTGATTTTGCTGGCTCTAATTCCCAAATGGCT
CAAGTCGGTGACGGTGATAATTCACCTTTAATGAATAATTTCCGTCAATATTTACCTTCCCTCCCTCAATCGGTTGAA
TGTCGCCCTTTTGTCTTTGGCGCTGGTAAACCATATGAATTTTCTATTGATTGTGACAAAATAAACTTATTCCGTGGT
GTCTTTGCGTTTCTTTTATATGTTGCCACCTTTATGTATGTATTTTCTACGTTTGCTAACATACTGCGTAATAAGGAG
TCTTAAAAGCTTGGGCCCGAACAAAAACTCATCTCAGAAGAGGATCTGAATAGCGCCGTCGACCATCATCATCATCAT
CATTGAGTTTAAACGGTCTCCAGCTTGGCTGTTTTGGCGGATGAGAGAAGATTTTCAGCCTGATACAGATTAAATCAG
AACGCAGAAGCGGTCTGATAAAACAGAATTTGCCTGGCGGCAGTAGCGCGGTGGTCCCACCTGACCCCATGCCGAACT
CAGAAGTGAAACGCCGTAGCGCCGATGGTAGTGTGGGGTCTCCCCATGCGAGAGTAGGGAACTGCCAGGCATCAAATA
AAACGAAAGGCTCAGTCGAAAGACTGGGCCTTTCGTTTTATCTGTTGTTTGTCGGTGAACGCTCTCCTGAGTAGGACA
AATCCGCCGGGAGCGGATTTGAACGTTGCGAAGCAACGGCCCGGAGGGTGGCGGGCAGGACGCCCGCCATAAACTGCC
AGGCATCAAATTAAGCAGAAGGCCATCCTGACGGATGGCCTTTTTGCGTTTCTACAAACTCTTTTTGTTTATTTTTCT
AAATACATTCAAATATGTATCCGCTCATGAGAC

**Supplemental Figures**

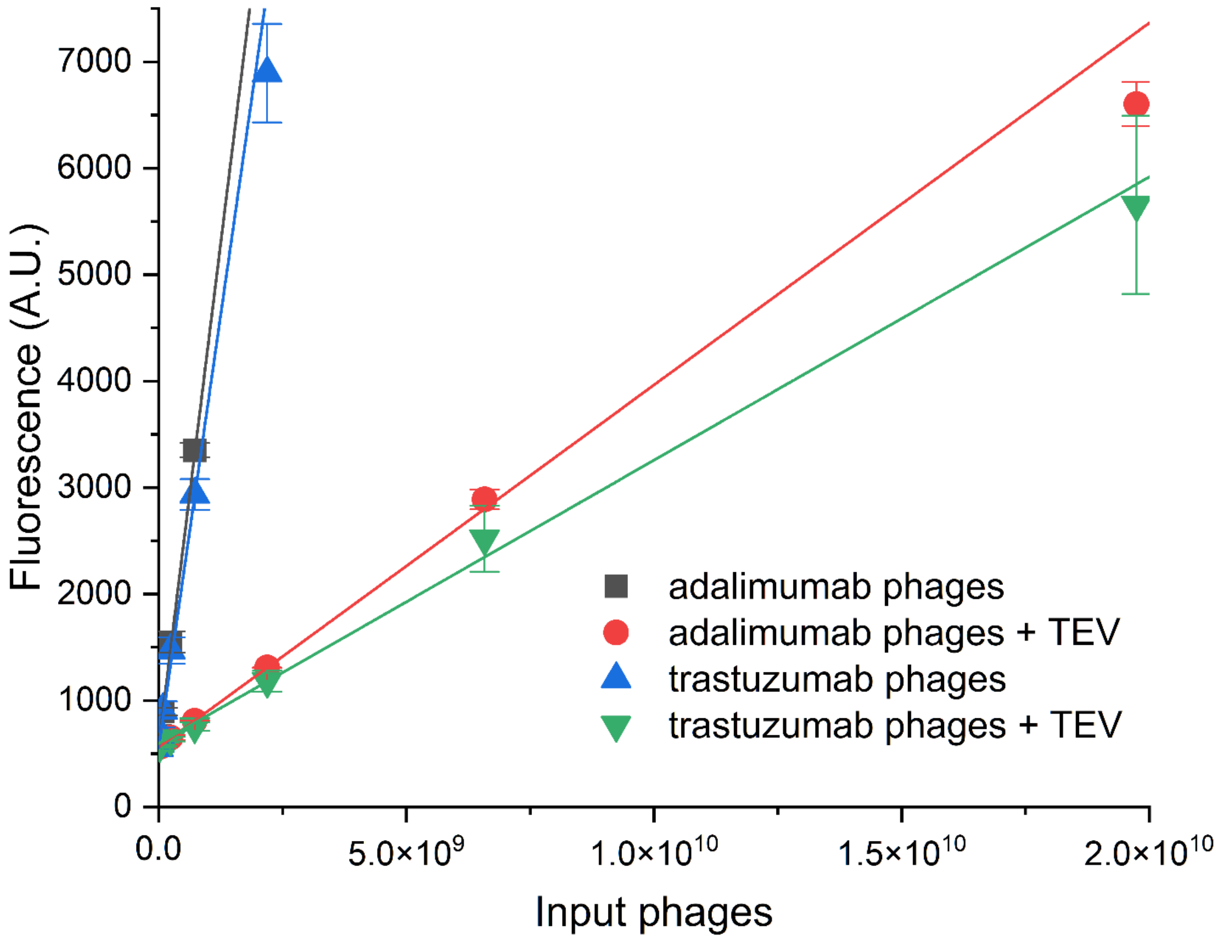


**Figure S1: Treatment of monovalent Fab phages in a dilution series with TEV protease**

Plotted are the linear parts and fits of anti-pVIII ELISAs quantifying the number of phages bound after 30 minutes of treatment with TEV or controls without TEV (Figure 3C). The background corrected signal of each curve at any phage number x can be derived from the linear fit equation: y-BG = m * x. At any given concentration x, the ratio of background corrected signals (y1-BG)/(y2-BG)=m1*x/(m2*x)=m1/m2. Dividing the slopes m of the TEV-treated fit with the control fit gives the ratio of remaining phages after treatment averaged over a range of concentrations and is the basis for figure 3C.

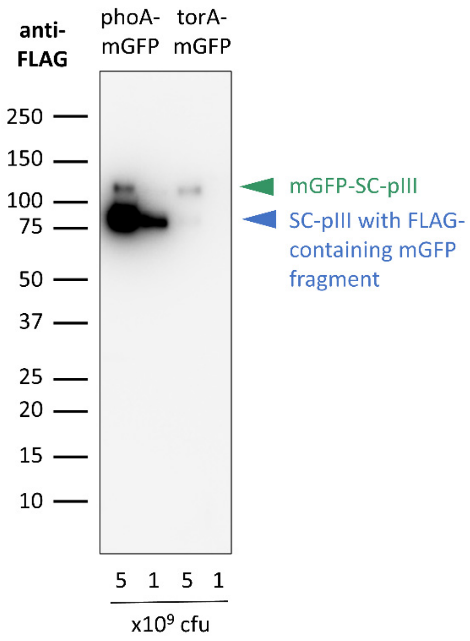


**Figure S2: Anti-FLAG immunoblot of displayed mGFP phages**

Immunoblot analysis of polyvalent SpyDisplay phages displaying mGFP-SpyTag with a PhoA or TorA leader peptide. Detection performed with anti-FLAG-HRP.

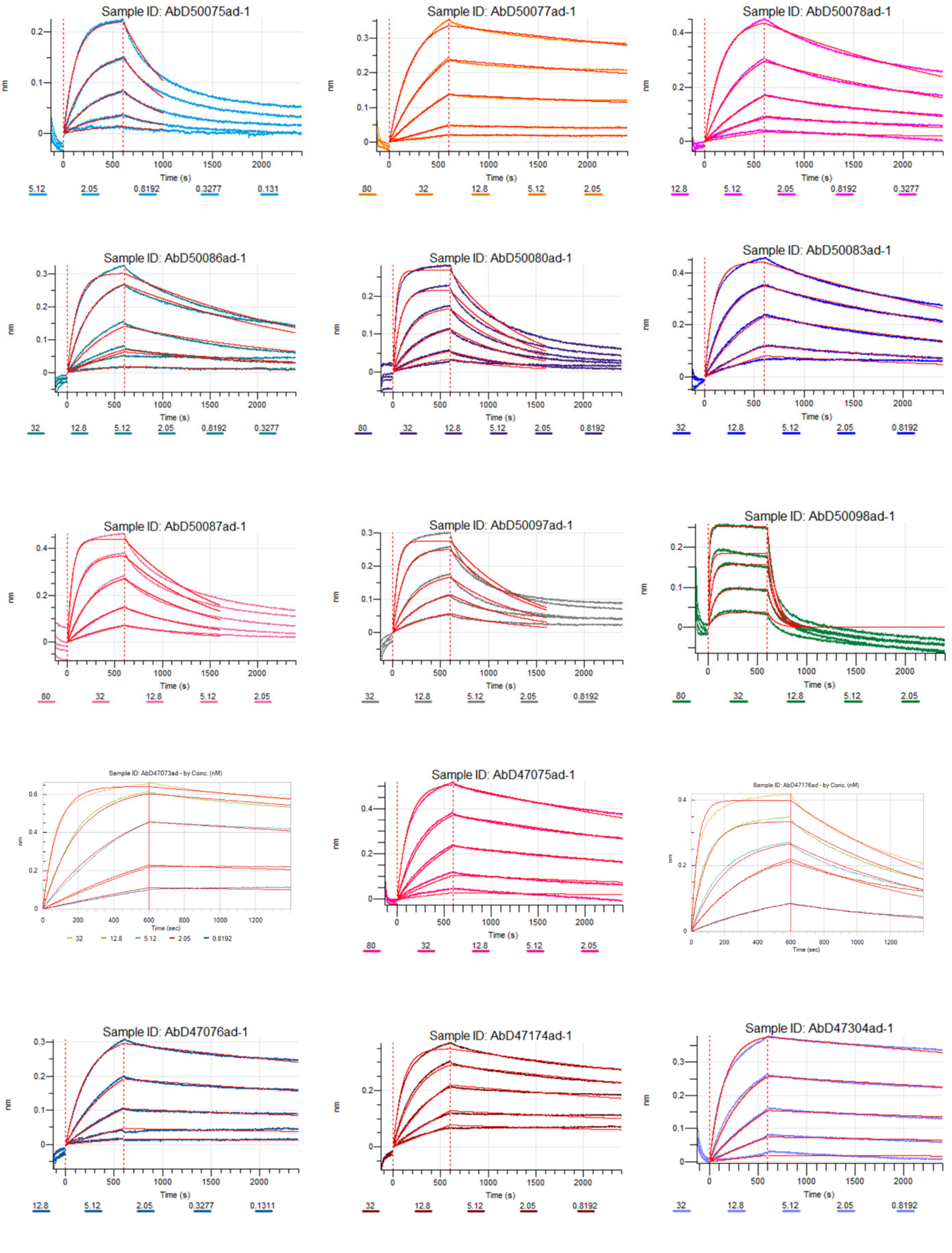


**Figure S3 A**: BLI sensorgrams of selected Fabs against mGFP

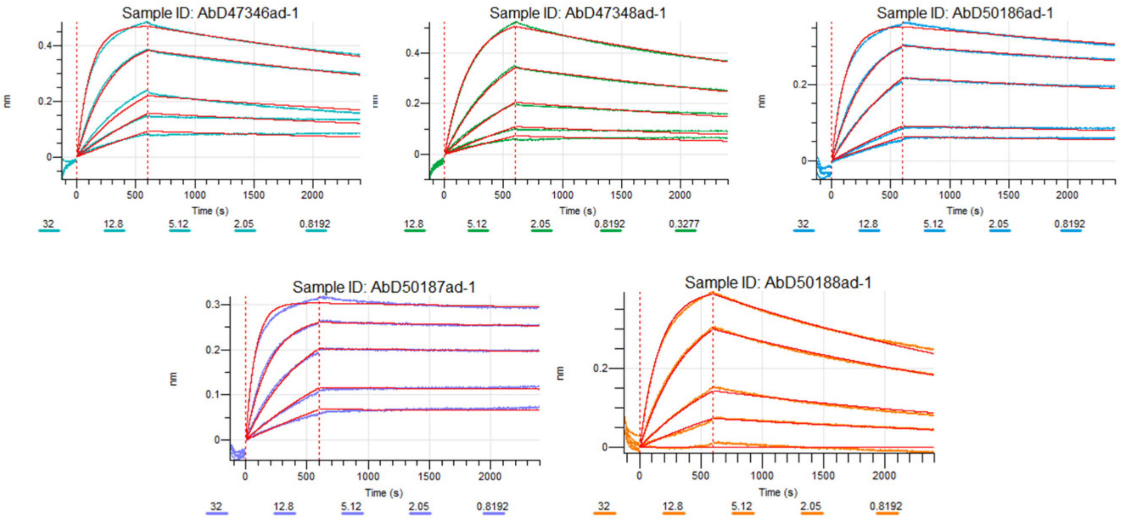


**Figure S3 A** (continued): BLI sensorgrams of selected Fabs against mGFP

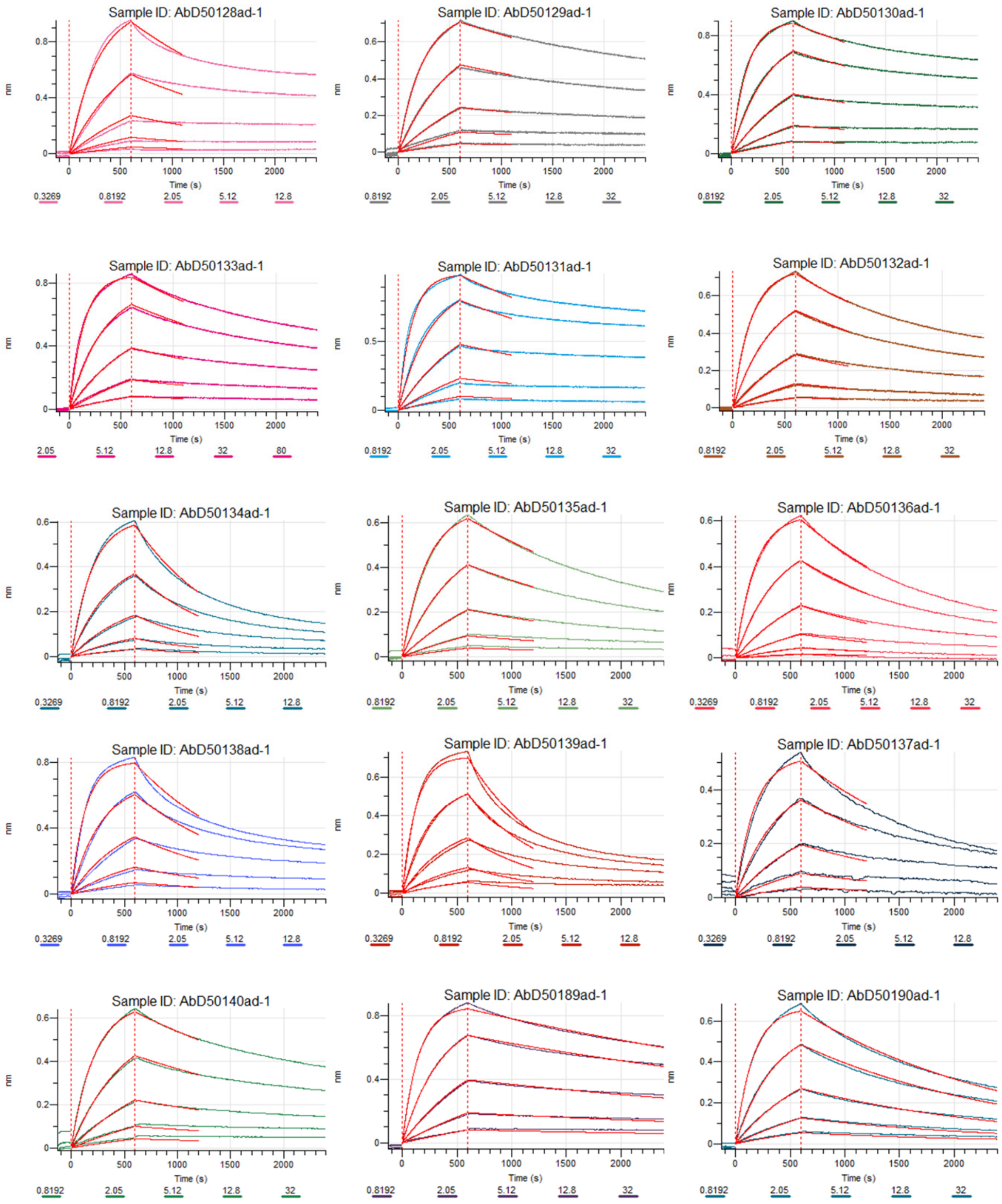


**Figure S3 B**: BLI sensorgrams of selected Fabs against sarilumab

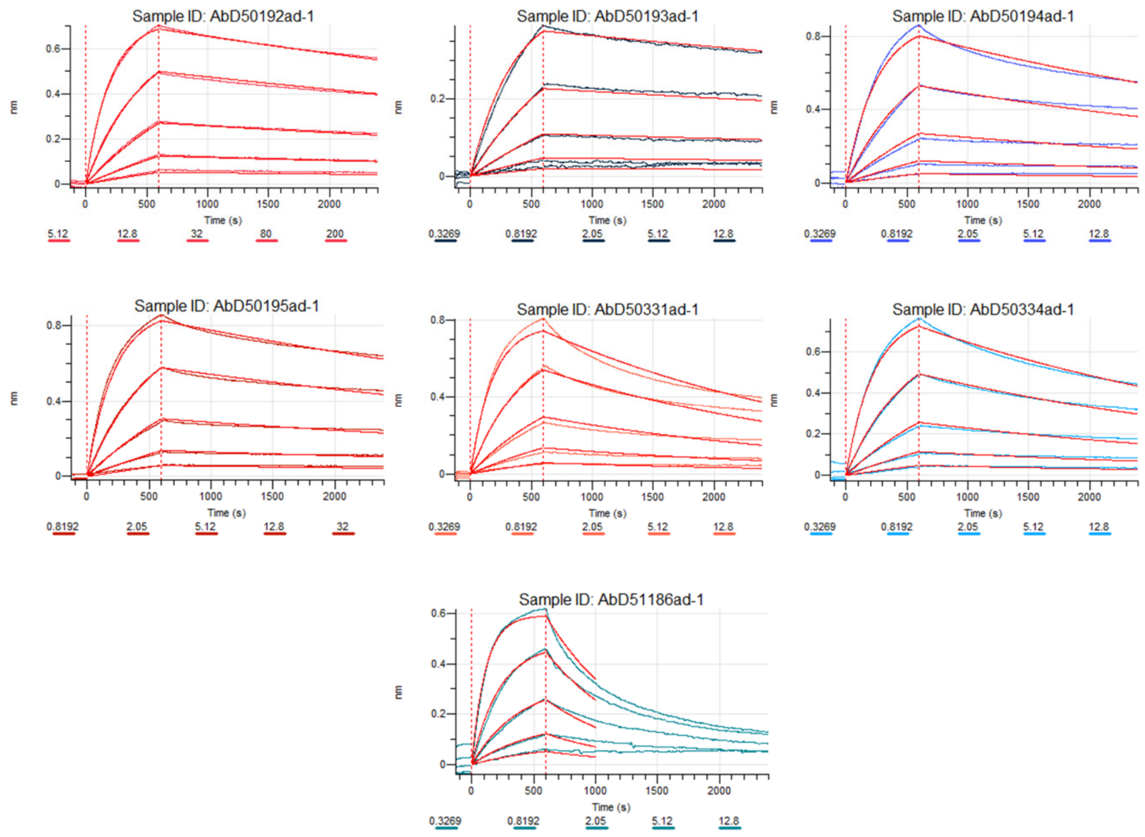


**Figure S3 B** (continued): BLI sensorgrams of selected Fabs against sarilumab

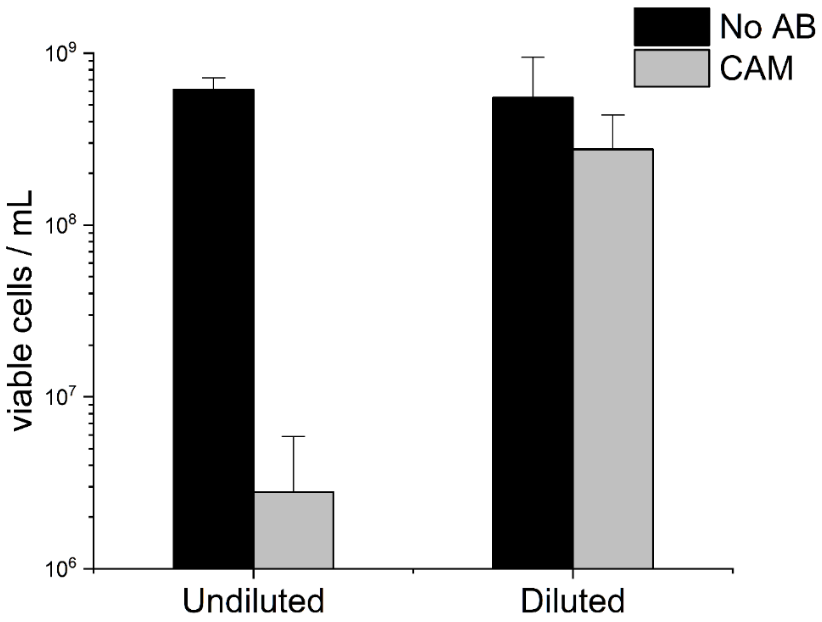


**Figure S4: Effect of dilution on viability of cells without phagemid in presence of chloramphenicol**

Concentration of viable SK25 bacteria in overnight cultures on chloramphenicol (CAM) or plates without antibiotic (No AB). Cultures were produced by infecting exponentially growing SK25 cells with Fab phages, incubating for 105 minutes, and adding 34 µg/mL CAM together with or without a 1:10 dilution step in fresh medium. Without a dilution step after phage rescue, uninfected cells are not selected against, and most cells do not carry the phagemid. The phenomenon of CAM not being effective in relatively dense bacterial cultures has been described (Guota, 1975) and might be related to bistability of bacterial growth depending on the level of expression of the resistance gene (Deris et al., 2013).
